# Episodic positive selection accumulates through time in seasonal influenza A, with no climatic signature

**DOI:** 10.1101/2025.11.20.689475

**Authors:** Matthieu Vilain, Rasha Mghabghab, Stéphane Aris-Brosou

## Abstract

The haemagglutinin (HA) and neuraminidase (NA) genes of seasonal influenza A evolve under continual immune-driven positive selection. To test whether the tempo of selection has changed over time, we mapped branch- and site-specific episodic diversifying selection (MEME) onto Bayesian relaxed-clock time trees for HA and NA in H1N1 and H3N2, across multiple countries and four sequence-subsampling schemes. We dated each selection episode and tested whether episodes accumulated through time after accounting for the growing number of sampled lineages. Positive-selection episodes increased over time in every gene-subtype combination, at about 2-6% per lineage-year, and rose faster for NA than HA. Episodes were concentrated at a small number of codon sites, especially recurrent sites in H3N2 HA that fell within canonical antigenic regions of the HA1 head. This increase was robust to subsampling scheme and time-bin width, and was driven disproportionately by recent lineages. A detrended spatial analysis found no association with latitude or temperature anomalies. Overall, positive selection on influenza surface antigens appears to be intensifying through time, most likely because of immune escape and expanded surveillance rather than climate warming.

## 1. Introduction

Seasonal influenza A viruses cause an estimated 290,000-650,000 respiratory deaths each year [1], and persist despite widespread vaccination because they continually escape host immunity through antigenic drift [2,3]. The surface glycoproteins haemagglutinin (HA) and neuraminidase (NA) are the principal targets of this process: positively selected substitutions accumulate in their antibody-binding epitopes, allowing successive variants to evade immunity acquired through prior infection and vaccination [4,5]. Characterising where and when this positive selection acts is therefore central to understanding, and anticipating, influenza evolution.

Episodic, lineage- and site-specific positive selection is now routinely inferred from sequence alignments [6,7], and influenza adaptation is known to proceed in punctuated bursts rather than at a steady pace [8], with appreciable genome-wide rates of adaptive substitution [9]. Yet these analyses almost always summarise selection over an entire phylogeny, yielding a single, static catalogue of selected sites or branches. Whether the *rate* at which episodes of positive selection occur has itself changed over the many decades that H1N1 and H3N2 have circulated, and whether any such change reflects intrinsic evolutionary dynamics or external factors such as a warming climate, has, to our knowledge, never been tested.

Answering this outstanding question requires placing individual selection events on an absolute time axis. This can be achieved through a two-step process, where (i) time-calibrated genealogies are estimated under relaxed molecular clocks [10], which underpins modern viral phylodynamics [11,12]; and (ii) combining them with branch-level inferences of selection, hence allowing each episode of positive selection to be dated and the temporal density of such episodes to be modelled. Here we integrate site- and branch-specific tests of episodic diversifying selection with Bayesian time trees for the HA and NA genes of the H1N1 and H3N2 subtypes, across multiple countries and four sequence-subsampling schemes. We ask (i) whether episodes of positive selection accumulate through time once the increasing number of sampled lineages is accounted for, (ii) whether the trend differs between genes and subtypes, and finally (iii) whether this trend carries any spatial signature of local climate and/or climate warming.

## 2. Material and methods

### (a) Sequence data and subsampling

Complete coding sequences of the haemagglutinin (HA) and neuraminidase (NA) genes of influenza A virus subtypes H1N1 and H3N2 were obtained from GenBank and the GISAID EpiFlu database for twelve countries (Brazil, Canada, Chile, China, India, Italy, Japan, the Netherlands, Russia, Singapore, Thailand and the USA; ESM). For each gene-subtype-country combination, sequences were aligned as in-frame codon alignments with MAFFT v7.526 [13]. Because the resulting alignments were very large and dominated by near-identical sequences, we reduced each to a tractable, representative subset using two complementary schemes. First, sequences were clustered with CD-HIT v4.8.1 [14,15] at two nucleotide-identity thresholds, 99% and 99.7%, retaining one representative per cluster. The more permissive 99% threshold yielded 29-293 sequences per alignment, while the stricter 99.7% threshold up to ∼2700.

Second, to distinguish the effect of phylogenetic thinning from that of the particular representatives chosen by clustering, we drew random subsamples without replacement, matched to the number of sequences retained by CD-HIT for each alignment and threshold. This produced four parallel sets of alignments (CD-HIT-99%, CD-HIT-99.7%, random-99% and random-99.7%) spanning the same gene-subtype-country combinations, with sequence collection dates ranging from 1933 to 2023.

### (b) Detection of episodic positive selection

For every alignment, each codon was tested for episodic diversifying (positive) selection with the mixed-effects model of evolution (MEME) [6], as implemented in HyPhy v2.5.78 [7]. For this, a maximum-likelihood tree was first inferred (SEM §1). MEME estimates a site-wise synonymous rate and a two-category mixture of non-synonymous rates, allowing a fraction of branches at a site to evolve under positive selection (*ω* > 1). We considered a codon to be under episodic selection at *p* ≤ 0.05 and, at such sites, attributed the signal to individual lineages using the empirical Bayes factor (*EBF*) for the positively selected rate class, flagging a branch when its *EBF* exceeded 100. Each branch × site combination satisfying both criteria was recorded as a discrete selection episode (SEM §2).

### (c) Time-scaled phylogenies

For each alignment, a time-scaled genealogy was inferred in BEAST v10.5.0 [10] under an HKY+Γ nucleotide substitution model [16] and an uncorrelated lognormal relaxed molecular clock [17], with a constant-size coalescent tree prior; tips were calibrated by their collection dates. Markov chain Monte Carlo chains were run for 10^8^ generations, and sampled every 5,000, discarding the first 50% as burn-in; convergence and effective sample sizes (> 200) were verified in Tracer v1.7 [18]. For each dataset we retained the maximum-a-posteriori (MAP) tree from the posterior distribution.

### (d) Dating selection episodes and testing for a temporal trend

Selection episodes were placed onto the corresponding BEAST MAP tree. As MEME and BEAST were run independently, branches were matched between the maximum-likelihood and Bayesian trees by their descendant taxa, terminal branches by accession and internal branches by their bipartition. Approximately 95-98% of episodes per dataset mapped to a BEAST branch, the remainder lying on internal branches that differed between the two topologies. Node calendar dates were obtained from the branch-length structure of each MAP tree, anchored at the most recent collection date, and each episode was dated at the midpoint of its branch (SEM §3).

To test whether episodes accumulate through time while accounting for the increasing number of lineages towards the present, episodes were binned into 3-year intervals. For each interval, we computed the available phylogenetic exposure as the total length (in lineage-years) of the time tree overlapping that interval. For each dataset, we fitted a Poisson generalised linear model of episode counts on calendar year with log(exposure) as an offset (the branch-length-normalized rate) and, for comparison, without the offset (raw counts), using statsmodels in Python 3 [19]. Per-dataset slopes were combined across datasets by inverse-variance random-effects meta-analysis [20] and by a sign test. A pooled Poisson model was then used, with a per-dataset intercept and year × gene and year × subtype interactions, to test whether the temporal trend differed between HA and NA and between H1N1 and H3N2 (ESM §4). Results were robust to bin width (2-5 years) and to restricting episodes to internal branches (ESM §5). Datasets whose label collection dates were inconsistent with the inferred BEAST time scale (one HA dataset) were excluded from the temporal analyses.

## 3. Results

Across the four sampling schemes we recovered, 4,595 branch × site episodes of episodic positive selection (*p* ≤ 0.05, *EBF* > 100), of which 94.6-98.4% could be placed and dated on the corresponding BEAST trees. The few unmatched episodes laid on internal branches that differed between the maximum-likelihood and Bayesian topologies. After normalizing for the number of lineages available through time, the rate of episodic positive selection was found to increase with calendar time in every scheme (random-effects meta-analysis of per-dataset slopes: all *p* ≤ 3.1 × 10^−5^; 12-26 of 16-27 datasets per scheme with positive slopes; ESM §4-5).

The increase was pervasive across the influenza surface antigens: in all sixteen gene × subtype × scheme combinations the temporal slope was positive and statistically significant (figure 1b). Pooled within genes and subtypes, the episode rate rose by roughly 2-6% per lineage-year (HA, 1.9-3.4%; NA, 1.9-5.7%; H1N1, 2.3-5.7%; H3N2, 1.9-5.7%), producing the steep accumulation of episodes towards the present shown in figure 1a. The increase was consistently steeper for NA than for HA (year × gene interaction *p* = 0.032, 3.0 × 10^-5^ and 3.4 × 10^-3^ in the CD-HIT-99%, CD-HIT-99.7% and random-99.7% schemes, respectively; *p* = 0.51 in random-99%), whereas H1N1 and H3N2 increased at statistically indistinguishable rates in every scheme (year × subtype *p* ≥ 0.11).

**Figure 1.**
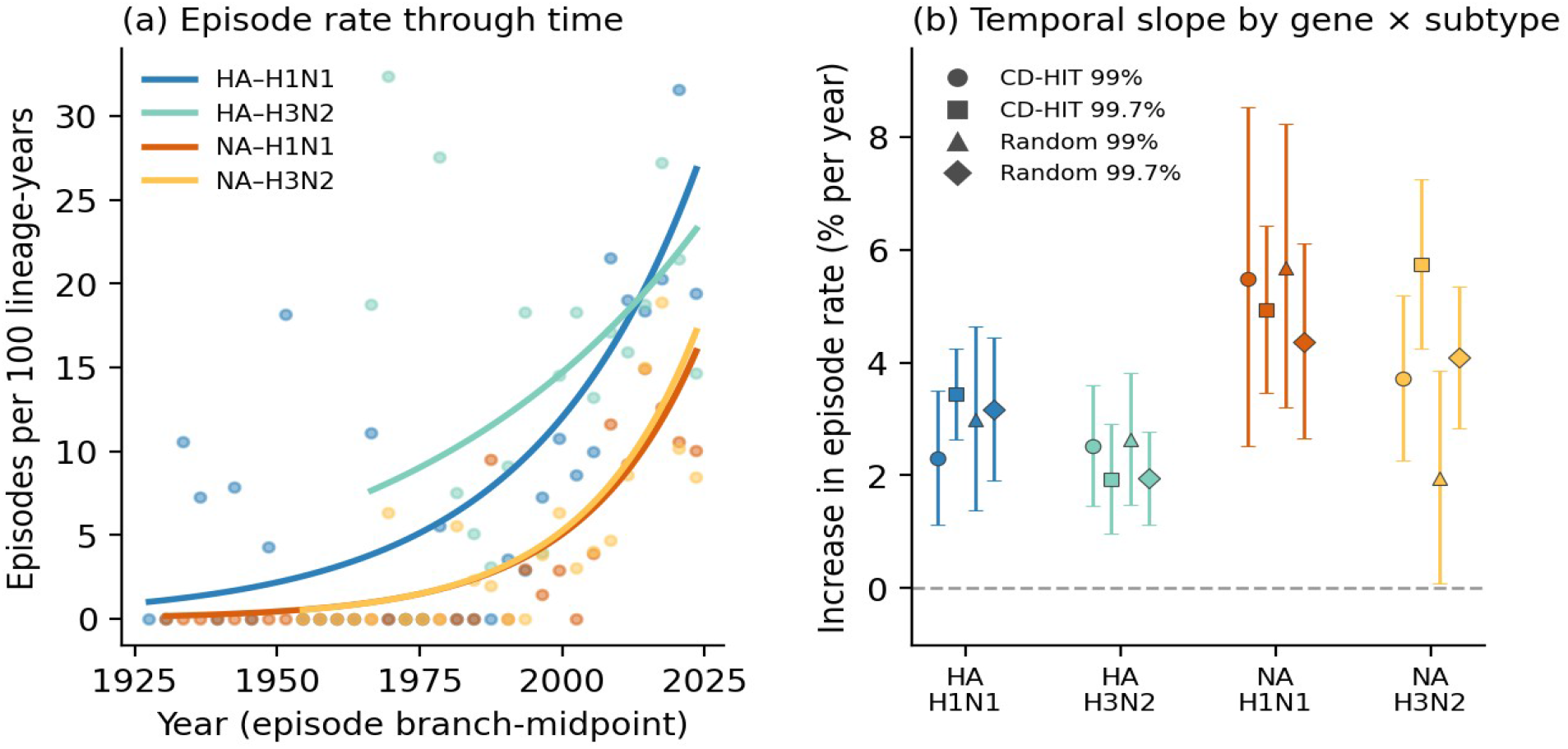
Episodic positive selection accumulates through time across influenza genes, subtypes and sampling schemes. (a) Rate of MEME-inferred selection episodes (episodes per 100 lineage-years, normalized by the available branch length) against the calendar date of each episode’s branch midpoint, for the four gene×subtype combinations in the CD-HIT-99.7% data; points are pooled 3-year bins and curves are fitted Poisson trends. (b) Estimated increase in episode rate (% per lineage-year, ±95% CI) for each gene×subtype combination under each of the four sampling schemes. All sixteen estimates are positive and significant.

The trend was robust to analytical and sampling choices. Estimates were essentially unchanged across 2-5-year time bins, and the four sampling schemes, two CD-HIT thresholds and size-matched random subsamples, gave congruent results (figure 1b), indicating that the signal reflects the data rather than the subsampling procedure (ESM §5). Raw, unnormalized episode counts increased about two-to three-fold faster than the branch-length-normalized rate, confirming that part of the apparent acceleration simply tracks the growth in the number of sampled lineages towards the present. Restricting episodes to internal branches, the increase remained significant in two schemes (CD-HIT-99% and random-99.7%), was marginal in random-99% (*p* = 0.07) and non-significant in the most densely sampled CD-HIT-99.7% set, indicating that the recent rise is carried disproportionately by terminal lineages. A country-level analysis using the observed CRU CY v4.08 year-by-year country temperatures found no association between the episode rate, or its temporal slope, and either country latitude, mean temperature, or the temperature anomaly detrended from each country’s warming trend, once sampling and calendar year were controlled (ESM §6).

## 4. Discussion

Across two surface antigens, two subtypes and four independent subsampling schemes, episodes of positive selection in seasonal influenza A accumulated through time, even after accounting for the increasing number of lineages sampled towards the present. This is consistent with the continual antigenic drift by which influenza escapes accumulating population immunity [2,5], and with earlier reports of recurrent, lineage-specific bursts of positive selection in the virus [4,8]. A rising rate of selection episodes per unit of evolutionary time is what one would expect if host-population immunity, built up over decades of infection and, increasingly, vaccination, impose progressively more frequent selective pressure on circulating viruses [2,9]. Consistent with this interpretation, the episodes are strongly concentrated on a small minority of codon sites in every gene×subtype combination (top ten sites carry 55-71% of episodes), and the most-recurrent sites in H3N2 HA are repeatedly hit by independent episodes across multiple countries, mapping to the canonical antigenic regions of the HA1 head (ESM §7).

The temporal increase was steeper for neuraminidase than for haemagglutinin in three of the four schemes. Although HA has long been the canonical target of antigenic drift, NA evolves under its own immune pressure and is increasingly recognised as an important, partially independent antigen [21]; the added selective pressure of NA-inhibitor antivirals deployed from the late 1990s may also contribute. The finding that NA selection has intensified at least as fast as HA selection, reinforces the case for monitoring NA evolution alongside HA’s. By contrast, H1N1 and H3N2 showed statistically indistinguishable temporal trends, suggesting the phenomenon is a general feature of seasonal influenza A rather than a peculiarity of one subtype.

An important limitation is that the apparent acceleration may partly reflect statistical power rather than biology. Tests for episodic selection gain power as branches accrue more substitutions and as clades are more densely sampled [22], conditions met more often in the recent, well-sampled portions of the genealogy. Normalizing by branch length removes the dependence on the number of lineages, but not this dependence on detection power. Two features of our results temper, without eliminating, this concern: the trend is remarkably consistent across genes, subtypes, identity thresholds and random versus clustered subsampling. Yet, it is carried disproportionately by terminal branches and, in the most densely sampled dataset, disappears on internal branches alone. We therefore regard the direction of the trend as robust, but interpret its magnitude conservatively.

Finally, because the temporal increase in the rate of episodic selection is collinear with calendar time, it cannot by itself be attributed to any particular external driver. We tested temperature directly using the observed year-by-year country temperatures of the CRU CY v4.08 dataset [23], decomposing each country’s series into a long-term warming trend and a detrended interannual anomaly: only the anomaly is statistically independent of calendar time and therefore identifiable as a temperature effect. The anomaly had no effect on the episode rate (*p* = 0.54), while calendar year retained its full +3.7% per-year increase in the same model (*p* = 4×10^−4^); countries that warmed faster did not accelerate faster (Spearman *ρ* = 0.31, *p* = 0.33), tropical and temperate countries showed indistinguishable episode rates and temporal slopes (Mann-Whitney *p* ≥ 0.57; ESM §6.5), and the only weak between-country tendency ran counter to a warming effect. Therefore, while climate may influence influenza *transmission* and seasonality, we find no positive evidence that a warming climate drives the accelerating selection we observe. The pattern is most parsimoniously explained by intensifying immune-driven escape compounded by changing sampling. While future work could integrate epidemiological relevant features such as R_0_ or case counts, we show here that mapping site- and branch-specific selection onto time-calibrated genealogies nonetheless places discrete adaptive events on an absolute time axis [12], and offers a general way to ask when selection acts [24].

## Supporting information

ESM

## Data accessibility

All scripts supporting the findings of this study are available from GitHub (https://github.com/sarisbro/data).

