## Supplementary material for "Episodic positive selection accumulates through time in seasonal influenza A, with no climatic signature": ESM

*Electronic Supplementary Material*

### 1. Supplementary notes on data preprocessing and evolutionary analyses

We paired two analyses for each influenza dataset: MEME [1] episodic-selection results (HyPhy v2.5 [2]) and a BEAST [3] relaxed-clock maximum-a-posteriori time tree. 30 datasets carry both files and were analysed, spanning genes HA and NA, subtypes H1N1 and H3N2, and 12 countries. Datasets present as only a tree or only a MEME run were not used.

Taxa are shared between the two analyses whilst formatted slightly differently (MEME: *AAO46463_CANADA_H3N2_NA_NONE_1968_01_01*; BEAST: *cds:AAO46463\Canada\H3N2\NA\none\1968_01_01*). The GenBank/EPI accession (e.g. AAO46463) is the stable key and was used to align the two trees. Tip sets matched exactly in every dataset checked.

#### 1.1 Preprocessing and analysis pipeline

The path from each BEAST input XML to the corresponding MEME analysis comprised eight scripted, parallelisable steps; all scripts are available from the project repository (https://github.com/sarisbro/data). The BEAST run itself, from which the time-calibrated MAP tree was retained, is described in §2 of the main text and reused unchanged here.

1. **Extract coding alignments from BEAUTi XML**. For every BEAUTi-generated XML file we parsed the <*alignment*> block and wrote the in-frame coding sequences and taxon labels to a per-dataset FASTA file. This is the same alignment used by the BEAST run from which the time tree was estimated.
2. **Clean FASTA headers**. The BEAST-style labels of the form *cds:ACC\Country\Subtype\Gene\role\YYYY_MM_DD* were converted to a FastTree- and HyPhy-compatible form by stripping the *cds:* prefix and replacing each backslash with an underscore, yielding *ACC_Country_Subtype_Gene_role_YYYY_MM_DD*. The original accession (e.g. AAO46463) was preserved as the stable cross-tree key.
3. **Build a maximum-likelihood phylogeny per alignment**. For each cleaned FASTA file we inferred an ML tree with FastTree [4] under the GTR [5] + Γ [6] model (*-nt -gtr -gamma*); these trees served as the input for MEME.

4. **Re-root each ML tree on the oldest sequence**. The sampling date (YYYY_MM_DD suffix) of every tip was extracted, the oldest tip(s) were identified, and the tree was re-rooted on this outgroup using *ape::root(..., resolve.root = TRUE)* [7], placing the deepest divergence at the earliest sample, consistent with the heterochronous calibration.
5. **Replace internal stop codons** (Biostrings [8]). The alignment was read codon by codon and any internal TAA, TAG or TGA codon was replaced with three gap characters, since HyPhy MEME requires an in-frame coding alignment without spurious stop codons.
6. **Resolve ambiguous nucleotides column-wise**. For every alignment column, ambiguity codes (any symbol outside A, C, G, T or gap) were replaced with the most frequent unambiguous nucleotide observed at that column, or with N when no unambiguous base was present, to avoid MEME failing on IUPAC ambiguity codes.
7. **Assemble HyPhy-compatible NEXUS files**. For each combination, the cleaned codon alignment (step 6) and the rerooted ML tree (step 4) were merged into a single NEXUS file containing matched DATA and TREES blocks, with taxon names sanitised to a common set.
8. **Generate and run MEME jobs**. For every NEXUS file a Bash launcher was generated that invokes *hyphy meme --alignment <nexus> --branches All --output <json>*, scanning all branches of the ML tree for episodic diversifying selection. The resulting per-site, per-branch JSON outputs are the inputs for the downstream analyses described in §2 (defining and dating selection episodes) and §4–6 (modelling).

#### 1.2 Sequence-count comparability of HA and NA

We checked whether the number of sequences available per country differs systematically between HA and NA. Pooling subtypes, paired Wilcoxon tests on per-country $\log_2$(HA/NA) ratios are not significant in any of the four sampling schemes ($p \geq 0.10$; figure S0). Stratifying by subtype, however, reveals a consistent ~20-30% excess of HA over NA sequences for H1N1 (mean $\log_2$ ratio +0.19 to +0.31, every country), whereas H3N2 sampling is balanced (mean $\log_2$ ratio ≈ 0). The H1N1 vs H3N2 difference itself is significant (Mann-Whitney $p$ = 0.017-0.033 in CD-HIT schemes; $p$ = 0.016 in random-99.7%).

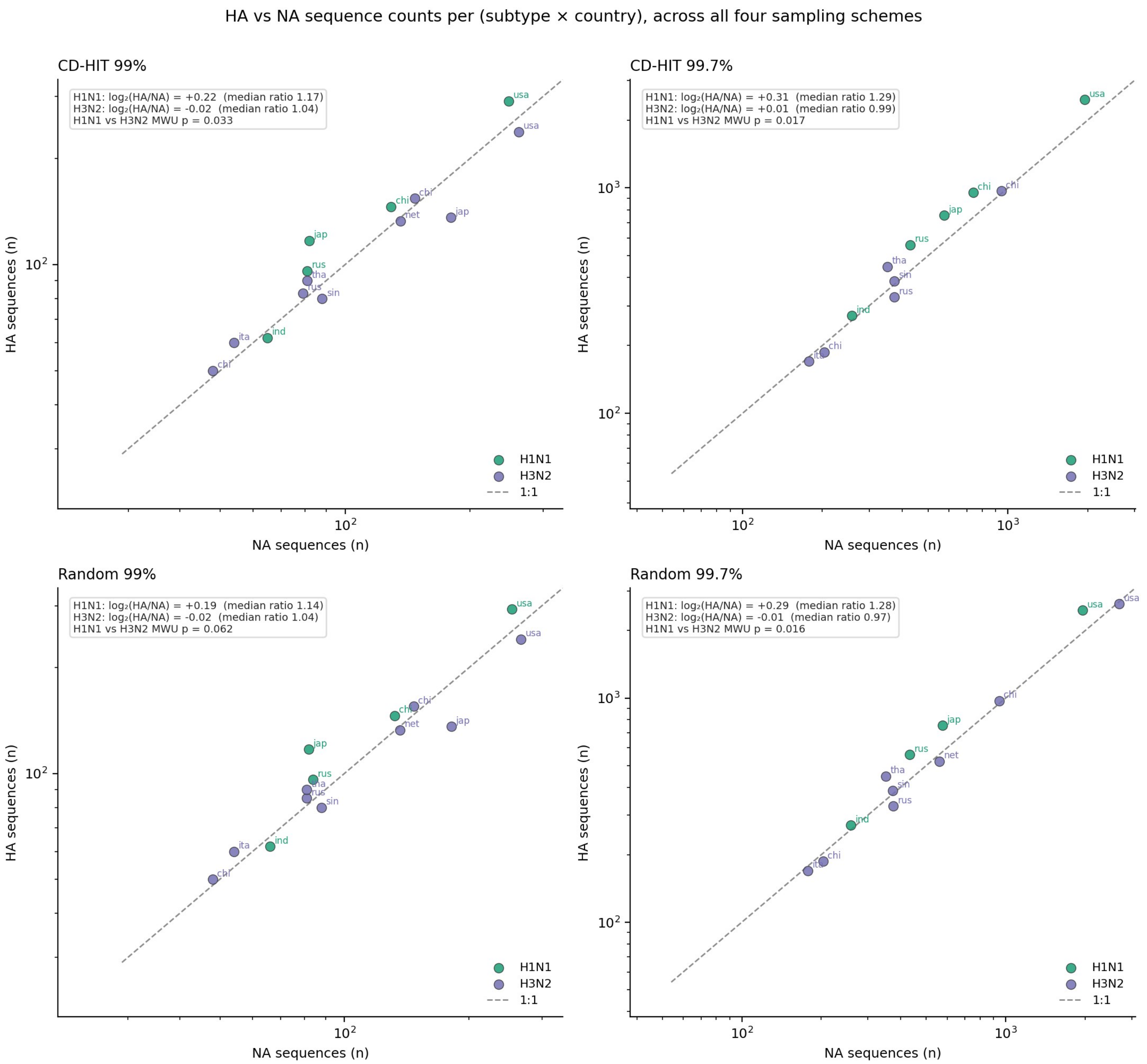


*Figure S0. HA and NA sequence counts per subtype vs country across all four subsampling schemes: (a) CD-HIT 99%; (b) CD-HIT 997.%; (c) Random 99%; and (d) Random 99.7%.*

This subtype-specific asymmetry plausibly reflects historical HA-prioritised surveillance for H1N1 since the 2009 pandemic. The temporal-rate models in r4 use log(n_tips) as a covariate and log(branch-length) as an offset, so this imbalance is partialled out; in particular, the H1N1 HA excess gives MEME more power to detect HA episodes than NA episodes for that subtype, yet the year × gene interaction (§4.3) still shows NA selection rising faster than HA.

### 2. Supplementary methods

#### 2.1 Defining an episode of positive selection

MEME reports, per codon site, a likelihood-ratio test for episodic diversifying selection and, per branch, the posterior probability that the branch belongs to the positively-selected rate class ($\beta^+$). For each site we computed the empirical Bayes factor (*EBF*) for every branch,

$$EBF = [\ Pr(\beta^+ \mid branch) / (1 - Pr(\beta^+ \mid branch))\ ] \div [\ p^+ / (1 - p^+)\ ]$$

where $p^+$ is the site's $\beta^+$ mixture weight (the prior). An *episode* is a (branch × site) pair at which the site is significant at MEME $p \leq 0.05$ and the branch has EBF > 100. This EBF construction was validated against MEME's own "# branches under selection" column, which it reproduced almost exactly (positive class = posterior-array row 1).

#### 2.2 Matching MEME branches to the BEAST time tree

MEME runs on a maximum-likelihood (ML) tree under GTR+Γ, rooted with the oldest sequence of each dataset, whereas dates come from the independently-estimated BEAST tree (also under GTR+Γ). Every MEME branch (terminal or internal) was matched to a BEAST branch by its bipartition (the set of descendant accessions). Terminal branches always match; internal branches match only where the two trees agree topologically.

#### 2.3 Dating an episode

Calendar dates were read off each BEAST tree from its own branch-length structure (root-to-node depth, anchored at the most-recent sampling date), which is robust to any mismatch between label dates and the inferred timescale and guarantees descendants are younger than ancestors. Each episode was timestamped at the midpoint of its matched BEAST branch (the mean of the parent-node and child-node dates), since MEME localises selection to a branch rather than a node.

#### 2.4 Testing for a trend through time

Episode timestamps were binned in calendar time. For each bin we also computed the available branch-length (the summed overlap of every BEAST branch interval with the bin, in lineage-years) as the exposure / opportunity for selection. We then fit, per dataset, a Poisson GLM of episode counts on calendar year:

- **Normalized model:** count ~ year, with offset log(branch-length), which tests whether episodes per lineage-year change through time.
- **Raw model:** count ~ year, no offset, which tests the unadjusted count trend (confounded by the growing number of lineages).

Per-dataset normalized slopes were combined by inverse-variance meta-analysis (fixed and DerSimonian–Laird random effects [9]), a sign test on slope direction, and a Stouffer combination of signed z-scores [10]. A pooled Poisson GLM across datasets (with a dataset fixed effect and *year*×*gene* and *year*×*subtype* interactions, branch-length offset), implemented in Python statsmodels [11], tested whether the temporal trend differs between HA/NA and H1N1/H3N2.

### 3. Matching and dating: coverage

Across the 30 datasets, 728 episodes (branch×site pairs) met the $p \leq 0.05$ & $EBF > 100$ criterion. Of these, 689 (94.6%) were matched to a BEAST branch and dated; the remaining 39 (5.4%) laid on internal branches absent from the BEAST tree (maximum-likelihood vs. BEAST topological discordance) and could not be dated. Unmatched episodes were concentrated in H3N2 NA Japan (30 episodes), where the ML and BEAST topologies disagree on many internal splits. Notably, 427 of 728 episodes (58.7%) fall on terminal branches.

Note that in one dataset (H3N2 HA Thailand), the sampling dates embedded in the tip labels disagree with the BEAST timescale (label-vs-tree deviation up to ~36 years). This tree collapses in calendar time and was excluded from all temporal tests.

### 4. Supplementary results: episodes increase through time

#### 4.1 Per-dataset trends

Among the 21 datasets with reliable dates and ≥ 8 dated episodes, 19 show a positive normalized slope and two negative (sign-test $p = 2.2\times10^{-4}$); nine are individually significant at $p < 0.05$. The forest plot shows the per-dataset slopes with the meta-analytic summaries (Fig. S1).

#### 4.2 Meta-analysis

The overall increase is highly significant (random-effects $p = 4.7\times10^{-8}$; between-dataset heterogeneity $I^2 \approx$ 38%; Table S1). Group-specific meta-analyses concur (HA $p = 4.8\times10^{-9}$; NA $p = 0.011$; H1N1 $p = 1.9\times10^{-5}$; H3N2 $p = 1.5\times10^{-4}$).

***Table S1.*** *Combined per-dataset normalized slopes.*

| Group | k | Slope/yr (RE) | p (random) | p (fixed) | +/− | Sign p | Stouffer Z | Stouffer p |
|---|---|---|---|---|---|---|---|---|
| ALL | 21 | +0.0274 | $4.7\times10^{-8}$ | $3.6\times10^{-13}$ | 19/21 | $2.2\times10^{-4}$ | 7.49 | $6.8\times10^{-14}$ |
| HA | 13 | +0.0231 | $4.8\times10^{-9}$ | $4.8\times10^{-9}$ | 13/13 | $2.4\times10^{-4}$ | 5.83 | $5.5\times10^{-9}$ |
| NA | 8 | +0.0387 | 0.011 | $8.6\times10^{-6}$ | 6/8 | 0.289 | 4.71 | $2.5\times10^{-6}$ |
| H1N1 | 7 | +0.0269 | $1.9\times10^{-5}$ | $5.1\times10^{-6}$ | 7/7 | 0.016 | 4.77 | $1.8\times10^{-6}$ |
| H3N2 | 14 | +0.0260 | $1.5\times10^{-4}$ | $1.5\times10^{-8}$ | 12/14 | 0.013 | 5.80 | $6.6\times10^{-9}$ |

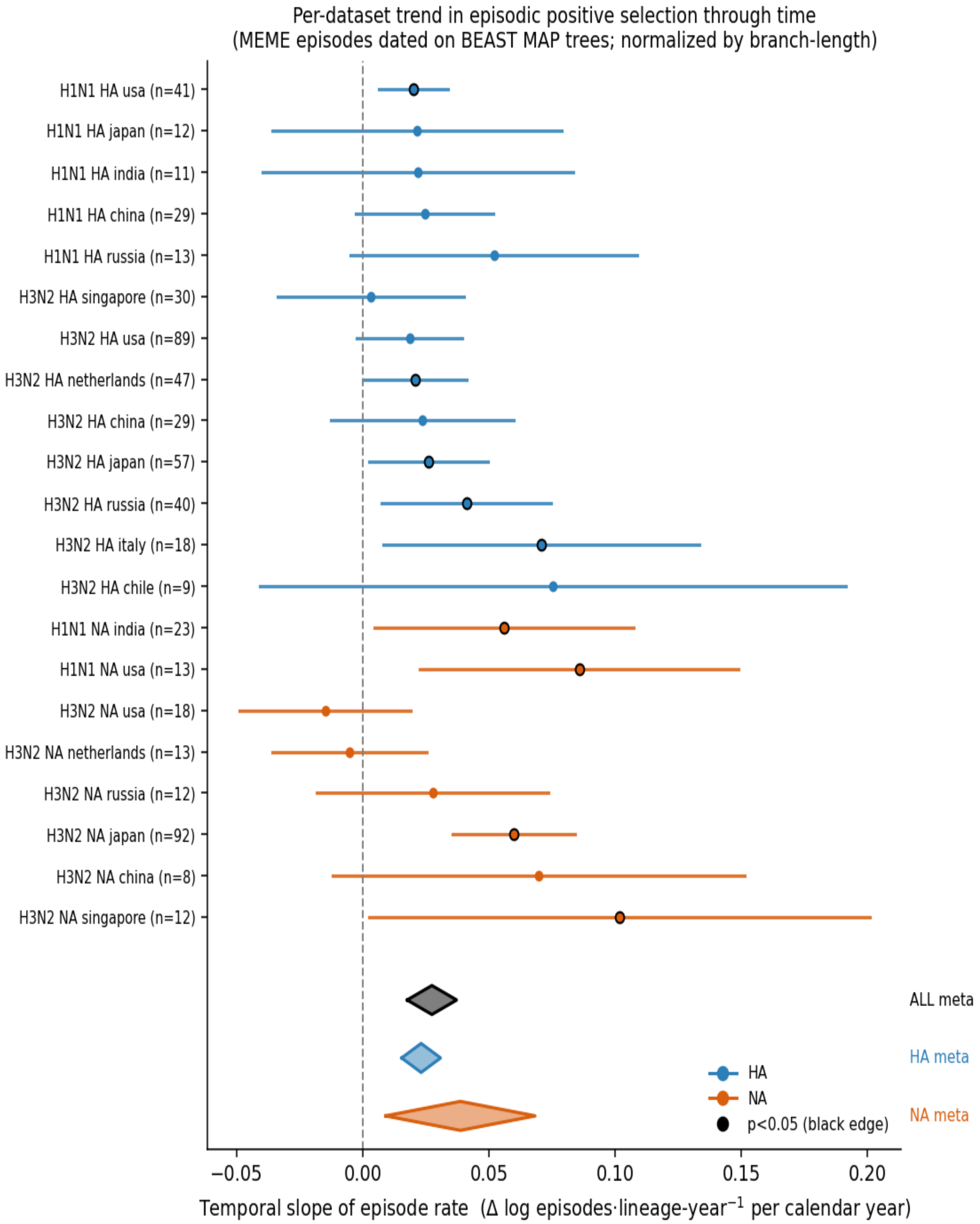


*Figure S1. Per-dataset slope of the normalized episode rate (Δ log episodes·lineage-year ⁻¹ per calendar year), 95% CI. Blue = HA, orange = NA; black-edged points are significant at p<0.05. Diamonds are random-effects meta-analytic summaries (ALL, HA, NA).*

### 4.3 Pooled rate through time, and HA vs NA / H1N1 vs H3N2

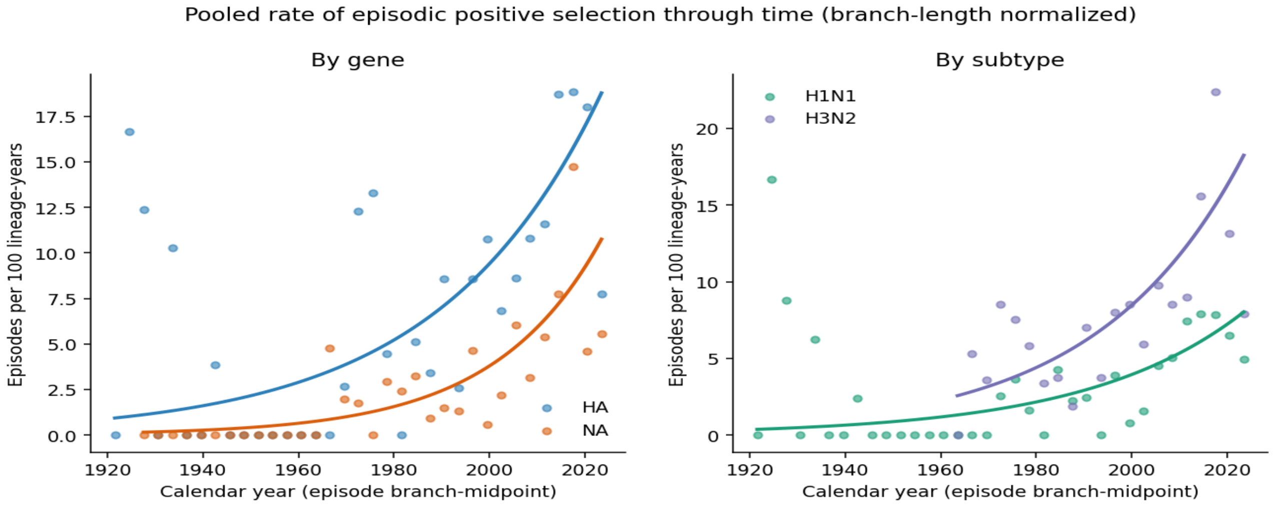


*Figure S2. Pooled episode rate (episodes per 100 lineage-years) versus calendar time, by gene (left) and subtype (right). Points are pooled 3-year bins; curves are fitted Poisson trends. Early-year points are noisy because few lineages (little branch-length) exist that far back.*

In a pooled Poisson GLM with a dataset fixed effect and branch-length offset, the calendar-year term is positive overall ($p = 1.3\times10^{-5}$; Fig. S2). Group-specific trends translate to roughly +2.4% per year for HA ($p = 1.7\times10^{-9}$) and +4.1% per year for NA ($p = 4.7\times10^{-10}$) in episodes per lineage-year. The year×gene interaction is significant ($p = 0.032$), showing that NA's episode rate rises faster than HA's. Subtype trends: H1N1 +2.9%/yr, H3N2 +3.0%/yr; the year×subtype interaction is not significant ($p = 0.793$), showing that the two subtypes increase at indistinguishable rates.

### 4.4 Why normalization matters

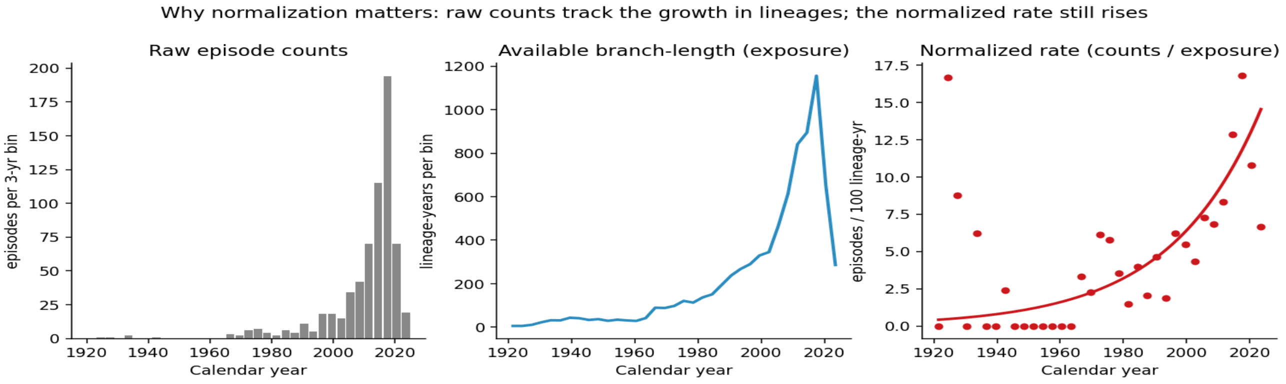


*Figure S3. Pooled raw episode counts (left) rise and then fall near the present, tracking the available branch-length / number of sampled lineages (middle). The normalized rate (right) removes this artifact and still increases monotonically, the genuine signal.*

Raw episode counts rise and then fall toward the present, mirroring the rise-and-fall of available branch-length (fewer lineages have reached the most recent years, and the final bin is partial; Fig. S3). Raw slopes are steeper than normalized slopes in 21/21 fitted datasets, confirming that lineage growth is a real confounding factor. The normalized rate therefore isolates the per-lineage increase, and must be taken into account.

### 5. Supplementary results: robustness

We assessed the robustness of the through-time increase in episodic positive selection with four sensitivity analyses, repeated independently for each of the four sequence-subsampling schemes. In each case the per-dataset Poisson slope of episode rate on calendar year was combined across datasets by inverse-variance meta-analysis; slopes are reported below as the percentage change in episode rate per year, $(e\beta-1)\times100$, with the meta-analytic $p$-value and the number of datasets with a positive slope. The four checks were: (i) varying the time-bin width (2, 3 and 5 years); (ii) restricting episodes to internal branches; (iii) restricting episodes to terminal branches; and (iv) replacing the branch-length-normalized rate with raw episode counts. Results are summarised in Fig. S4 and Tables S2–S4.

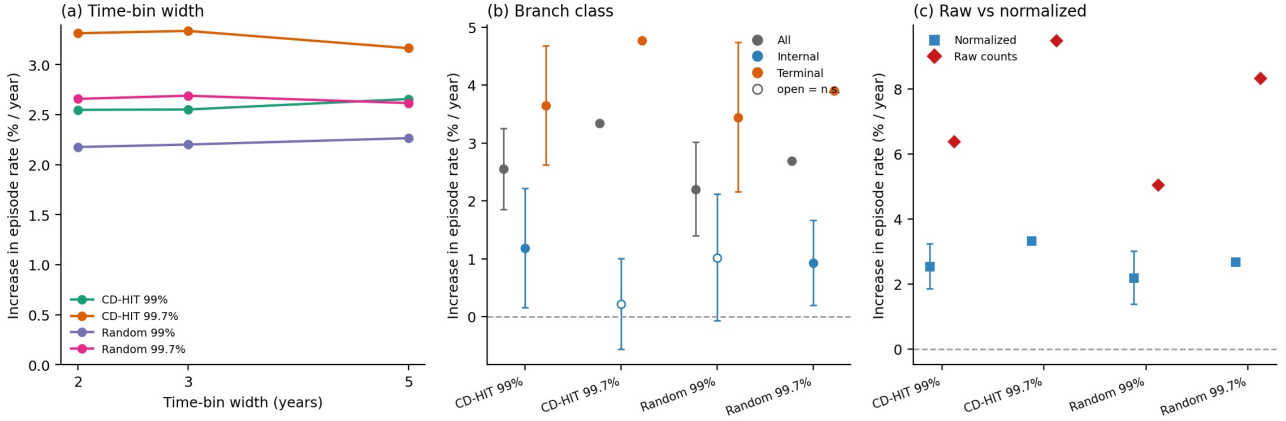


*Figure S4. Robustness of the temporal increase in episodic positive selection across the four sampling schemes. (a) Meta-analytic slope (% increase in episode rate per year) at 2-, 3- and 5-year time bins. (b) Slope for all episodes versus internal-branch-only and terminal-branch-only episodes (open symbols, not significant; bars are reconstructed 95% CIs). (c) Branch-length-normalized versus raw (unnormalized) episode-count slopes. The trend is insensitive to bin width, steeper on terminal branches and weak or absent on internal branches in the densest scheme, and inflated under raw counts.*

***Table S2.*** *Time-bin width: increase in episode rate (% per year) at three bin widths; the meta-analytic p-value (shown for the 3-year analysis) is* $< 10^{-7}$ *in every scheme. k = number of datasets combined.*

| Sampling scheme | k | 2-year | 3-year | 5-year | p (3-year) |
|---|---|---|---|---|---|
| CD-HIT 99% | 21 | +2.55 | +2.55 | +2.66 | $3.6\times10^{-13}$ |
| CD-HIT 99.7% | 27 | +3.31 | +3.34 | +3.17 | $2.2\times10^{-36}$ |
| Random 99% | 16 | +2.18 | +2.20 | +2.27 | $6.8\times10^{-8}$ |
| Random 99.7% | 24 | +2.66 | +2.69 | +2.62 | $7.9\times10^{-22}$ |

***Table S3.*** *Branch class: increase in episode rate (% per year) for all episodes and for episodes restricted to internal or terminal branches, with meta-analytic p and the number of datasets with a positive slope (positive/total).*

| Sampling scheme | Branch class | k | Slope (%/yr) | p | Datasets + |
|---|---|---|---|---|---|
| CD-HIT 99% | All | 21 | +2.55 | $3.6\times10^{-13}$ | 19/21 |
| CD-HIT 99% | Internal | 13 | +1.18 | 0.023 | 12/13 |
| CD-HIT 99% | Terminal | 19 | +3.65 | $1.3\times10^{-12}$ | 19/19 |
| CD-HIT 99.7% | All | 27 | +3.34 | $2.2\times10^{-36}$ | 26/27 |
| CD-HIT 99.7% | Internal | 19 | +0.22 | 0.582 | 11/19 |
| CD-HIT 99.7% | Terminal | 27 | +4.77 | $4.3\times10^{-43}$ | 26/27 |
| Random 99% | All | 16 | +2.20 | $6.8\times10^{-8}$ | 12/16 |
| Random 99% | Internal | 12 | +1.02 | 0.067 | 10/12 |
| Random 99% | Terminal | 14 | +3.44 | $9.7\times10^{-8}$ | 14/14 |
| Random 99.7% | All | 24 | +2.69 | $7.9\times10^{-22}$ | 23/24 |
| Random 99.7% | Internal | 21 | +0.93 | 0.013 | 15/21 |
| Random 99.7% | Terminal | 22 | +3.91 | $6.9\times10^{-21}$ | 22/22 |

***Table S4.*** *Raw versus normalized counts: branch-length-normalized rate versus raw episode counts (3-year bins). "Raw steeper" gives the number of fitted datasets in which the raw slope exceeded the normalized slope. The last column gives the percentage of episodes that could not be dated owing to maximum-likelihood/Bayesian topological disagreement.*

| Sampling scheme | Normalized (%/yr) | p | Raw (%/yr) | p | Raw steeper | Undated (%) |
|---|---|---|---|---|---|---|
| CD-HIT 99% | +2.55 | $3.6\times10^{-13}$ | +6.38 | $4.5\times10^{-87}$ | 21/21 | 5.4% |
| CD-HIT 99.7% | +3.34 | $2.2\times10^{-36}$ | +9.48 | $<10^{-300}$ | 27/27 | 4.3% |
| Random 99% | +2.20 | $6.8\times10^{-8}$ | +5.06 | $2.4\times10^{-39}$ | 16/16 | 1.6% |
| Random 99.7% | +2.69 | $7.9\times10^{-22}$ | +8.33 | $2.3\times10^{-215}$ | 24/24 | 5.1% |

The direction and approximate magnitude of the increase are stable across bin widths and across both clustered (CD-HIT) and size-matched random subsampling. The signal is steeper on terminal branches and weakest, or non-significant, on internal branches (notably in the most densely sampled CD-HIT-99.7% scheme). Raw counts rise about two- to three-fold faster than the normalized rate. Altogether, this indicates that the temporal increase is genuine, but is partly amplified by the growth of sampling and detection power toward the present.

- **Time-bin width.** Essentially unchanged across 2-, 3-, and 5-year bins (slope +0.0252, +0.0252, +0.0262/yr; all $p = 3.6\times10^{-13}$ or smaller; Table S2).
- **Internal branches only.** Excluding terminal branches, the meta-analytic slope is +0.0118/yr ($p = 0.023$; 12/13 datasets positive; Table S3). The increase is therefore not just an artifact of terminal branches being recent by construction; internal lineages show it too.
- **Terminal branches only.** +0.0359/yr, $p = 1.3\times10^{-12}$ (19/19 positive; Table S3).
- **Raw vs normalized.** Both increase; the raw-count slope (+0.0618/yr) is steeper than the branch-length-normalized slope (+0.0252/yr), which is the conservative estimate (Table S4).

### 6. Supplementary results: episode rate and its temporal trend show no association with temperature

The main analysis shows that episodes of positive selection accumulate through calendar time. Because global mean temperature has also risen monotonically over the study period, it, like any monotonic covariate of the period, including sequencing effort, is collinear with calendar year and cannot be tested as an independent cause from a single global time series. We therefore tested temperature using the observed year-by-year country temperatures of the Climatic Research Unit CY v4.08 dataset [12] and, critically, separated each series into its long-term warming trend and its detrended interannual anomaly: the anomaly is the part of temperature that is *not* a function of time, and is therefore the only part that can test temperature independently of the temporal trend itself.

#### 6.1 Decomposing temperature into a warming trend and a detrended anomaly

For each country $c$ we took the CRU CY v4.08 annual mean temperature series $T^{obs}(c, y)$ for years $y$ = 1933-2023 [12] and fitted a single straight line by ordinary least squares (OLS),

$$T^{fit}(c, y) = \alpha_c + \beta_c \cdot y,$$

where $\beta_c$ is the country's long-term warming rate (e.g. Russia β = +0.024 °C/yr, Japan +0.022, Chile +0.007). The temperature **anomaly** is the residual,

$$T^{anom}(c, y) = T^{obs}(c, y) - T^{fit}(c, y).$$

For a 3-year bin, we average over the bin's years. At every point in time, the observed temperature is thus split into two pieces : a smoothly increasing trend ,and a wiggle around that trend. The wiggle is the anomaly. Figure S5a makes this concrete for Japan: the red line is the fitted warming trend; the blue stems are the anomalies (residuals from the trend).

The point of the decomposition is a statistical orthogonality property. By construction of OLS residuals, the anomaly satisfies $\Sigma y\ T^{anom}(c, y) \cdot (y - \bar{y}) = 0$ for each country: within every country the anomaly has exactly zero linear correlation with calendar year. The trend component, by contrast, is a linear function of year (correlation 1). The decomposition therefore splits observed temperature into a piece that is perfectly collinear with time, and a piece that is uncorrelated with time. This is what allows temperature to be tested independently of the temporal trend itself.

#### 6.2 Models

For every dataset (gene × subtype × country × sampling scheme; $n = 113$), we computed the branch-length-normalized episode rate (matched episodes per lineage-year), the number of sequences (sampling effort), where estimable, the per-dataset temporal slope from the main analysis, and the exposure-weighted mean year. This last variable is the temporal centre of mass of the available phylogeny: a weighted average of calendar-bin midpoints, with each bin weighted by how much branch-length lived through that bin. It is intended to address the question as to "*when does this dataset sit,*" that is the period that the tree actually covers, weighted by how many lineages there are at each time: $\Sigma_i\ t_i \cdot E_i / \Sigma_i\ E_i$, where $t_i$ is the midpoint of calendar bin $i$ (3-year bins on a common grid), and $E_i$ is the exposure of bin $i$, i.e. the total branch length, in lineage-years, of the BEAST time tree overlapping that bin.

We then modelled episode counts with a Poisson generalized linear model (offset = log lineage-years; covariates log(number of sequences), gene, subtype and scheme; country-clustered robust standard errors) under three specifications:

- **A. Observed temperature only:** mirrors a naive rate-vs-temperature test; collinear with time within country (within-country median Pearson $r = 0.91$; figure S5a).
- **B. Observed temperature, with calendar year added:** does temperature contribute anything beyond year?
- **C. Detrended anomaly, with calendar year added:** the identifiable test of temperature, separated from the temporal trend.

We also related the per-country episode rate and its temporal slope to the country mean temperature, and the per-country temporal slope to the country warming rate (Spearman correlations, $n = 12$ countries; non-pseudoreplicated).

#### 6.3 Results

Within each country, observed temperature is almost perfectly collinear with calendar year (median Pearson $r = 0.91$; figure S5a), so observed temperature cannot, on its own, distinguish a temperature effect from a temporal trend. Consistent with this, observed temperature does not predict the episode rate (model A,

+0.03% per °C, $p = 0.971$), and adds nothing once calendar year is included (model B, $p = 0.454$). Most tellingly, the detrended temperature anomaly has no effect (model C, $p = 0.543$), while calendar year retains a strong, significant increase in the same model (+3.71% per year, $p = 4.1\times10^{-4}$; figure S5b,c). Between countries, neither the episode rate nor its temporal slope is associated with the country mean temperature (Spearman $\rho = 0.10$, $p = 0.762$; and $\rho = 0.01$, $p = 0.983$), and countries that warmed faster did not accelerate faster (slope vs warming rate $\rho = 0.31$, $p = 0.331$). In short, temperature predicts the trend only insofar as it is the trend; its time-independent component carries no signal.

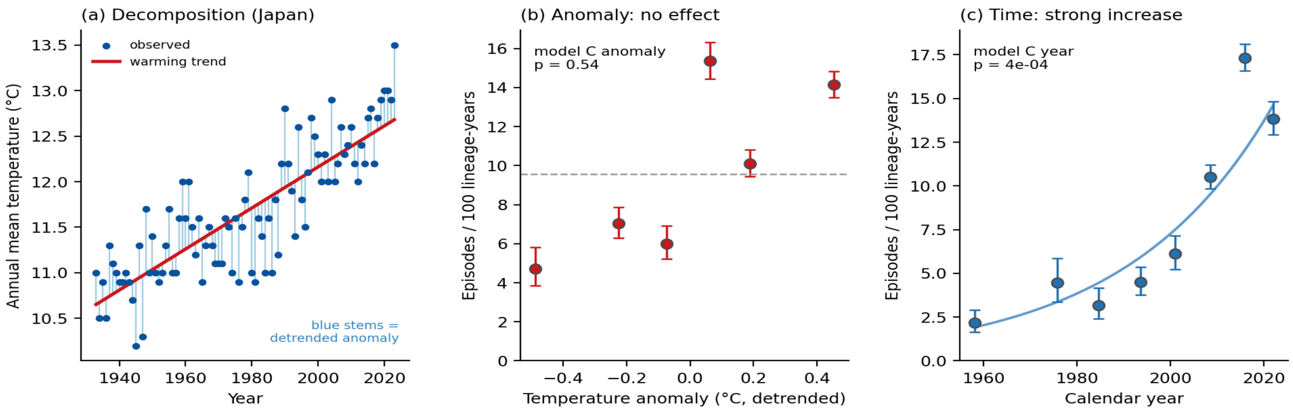


*Figure S5. Episode rate tracks the time-trend component of temperature, not its detrended anomaly (CRU CY v4.08). (a) Temperature decomposition for a representative country (Japan): the observed annual mean temperature (points) is the sum of a long-term warming trend (red line) and a detrended interannual anomaly (blue stems). (b) Pooled, exposure-weighted episode rate (±95% Poisson confidence interval) across bins of the detrended temperature anomaly: no relationship (dashed line, overall mean; model C anomaly $p = 0.54$). (c) The same pooled rate across calendar-year bins rises strongly (model C year $p = 4\times10^{-4}$). Temperature predicts the episode rate only through its time-trend component, which is calendar time.*

***Table S5.*** *Tests of an association between episodic positive selection and CRU CY v4.08 temperature. Rate models are Poisson with offset = log lineage-years and country-clustered SE; effects are % change in episode rate per unit predictor.*

| Test | Effect / statistic | *p* |
|---|---|---|
| Collinearity: temperature vs year (within-country) | $r = 0.91$ (median) | — |
| A. Observed temperature (per °C) | +0.03% | 0.971 |
| B. Observed temperature, year controlled (per °C) | −0.45% | 0.454 |
| C. Temperature anomaly, year controlled (per °C) | −23.14% | 0.543 |
| C. Calendar year, same model (per year) | +3.71% | $4.1\times10^{-4}$ |
| Between-country: rate vs mean temperature | $\rho = 0.10$ | 0.762 |
| Between-country: temporal slope vs mean temperature | $\rho = 0.01$ | 0.983 |
| Warming-rate: temporal slope vs warming rate | $\rho = 0.31$ | 0.331 |

Two limitations bound this inference: power is limited with twelve countries, and seasonal influenza circulates as a single, globally mixing population, so a country label largely records where a globally distributed lineage was sampled rather than a closed local selective environment. The spatial test can thus exclude a strong local-climate signature, but cannot identify a spatially uniform, purely temporal climate effect that remains confounded with calendar year. The result is consistent with the interpretation favoured in the main text: the temporal increase reflects rising detection power together with immune-driven antigenic escape, with temperature playing no detectable role.

#### 6.4 Per-country, per-gene × subtype expansion

Figure S6 expands figure S5 to every country and every gene × subtype combination. Each country panel shows the CRU CY v4.08 annual mean temperature (grey points) and its fitted linear warming trend (red line; right axis), together with the branch-length-normalized episode rate (left axis) for every gene × subtype combination present in that country, with per-cell Poisson trends. The decomposition is therefore shown country by country; the model fits are shown country × gene × subtype, with the CD-HIT-99.7% scheme used as the richest of the four sampling schemes. The figure demonstrates that the through-time increase in the episode rate is essentially universal across countries and gene × subtype combinations, while the temperature trend rises essentially linearly in every country with no obvious country-level dependence of the rate on the temperature trace beyond what the time axis itself carries.

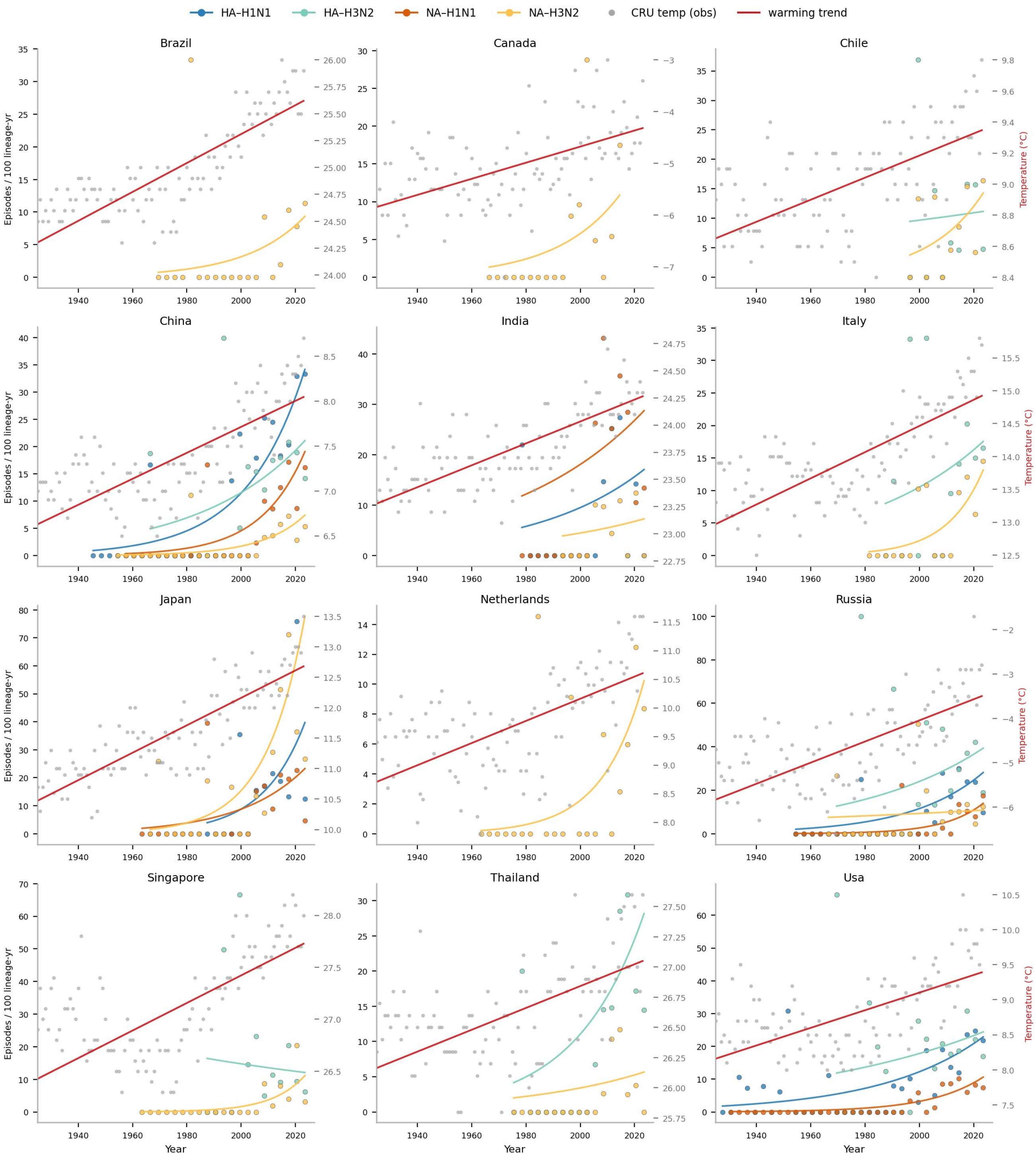


*Figure S6. Per-country temperature decomposition and per-country × gene × subtype episode-rate fits (CD-HIT-99.7% scheme). In each panel, grey points are the CRU CY v4.08 observed annual mean temperature and the red line is the fitted warming trend (°C, right axis). Coloured points and lines are the branch-length-normalised episode rate (episodes per 100 lineage-years, left axis) for each gene × subtype combination present in that country, with per-cell Poisson trends: HA–H1N1 (blue), HA–H3N2 (green), NA–H1N1 (orange), NA–H3N2 (yellow). Australia (no MEME data) and H3N2–HA Thailand (date-inconsistent; excluded from temporal tests) are not shown. Episode rates rise in almost every cell while temperature rises smoothly in every country; the rate trajectories do not track the temperature*

*traces beyond the shared monotonic increase with calendar time, consistent with the formal anomaly-detrended test in Figure S5 and Table S5.*

#### 6.5 Tropical versus temperate

Although the continuous tests in §6.1-§6.3 are the principal climate analysis, we also report the coarser tropical-versus-temperate contrast, given that influenza seasonality differs substantially. Countries were classed geographically by absolute centroid latitude (tropical = $|lat| < 23.5°$: Brazil, India, Singapore, Thailand; temperate = the remaining eight: Canada, Chile, China, Italy, Japan, the Netherlands, Russia, USA). For each dataset (gene × subtype × country × scheme) we computed the pooled episode rate per lineage-year and, where estimable, the per-dataset normalised temporal slope from the main analysis. We compared zones with Mann-Whitney U tests on the per-dataset rates and slopes, with a Poisson GLM of episode counts on zone (offset = log lineage-years; covariates log(sequences), calendar year, gene, subtype and scheme; country-clustered robust standard errors), and with a temperature anomaly × zone interaction in the same Poisson GLM to test whether the (overall null) detrended-anomaly effect differs between zones.

The two zones differ in observed CRU temperature by ≈20 °C as expected (tropical bin-level mean 26.0 °C versus temperate 6.6 °C; figure S5.5a). Despite this, the per-dataset episode rate and the temporal slope are statistically indistinguishable between zones: tropical and temperate medians of 13.13% and 14.49% per 100 lineage-years (Mann-Whitney $p = 0.62$; figure S5.5b), and slopes of 3.51% and 3.10% per year ($p = 0.57$; figure S5.5c). In the Poisson rate model, the tropical-zone coefficient is marginal and in the cold-skewed direction ($\beta = -0.27$, rate × 0.77 per °C-warmer zone, $p = 0.07$); the bulk-rate conclusion is therefore that climate zone, like absolute latitude and mean temperature in §6.3, does not predict the episode rate after sampling and time are controlled.

One nominally significant interaction did emerge: the detrended temperature anomaly × zone term was positive and significant ($\beta = 1.40$ per °C, p = 0.002), suggesting that within-country anomalies might predict episode rate more strongly in tropical than in temperate countries while the main anomaly effect remains null in temperate zones. We interpret this with strong caution: the tropical panel comprises only four countries ($n = 4$ clusters for the country-clustered standard error, well below the rule-of-thumb threshold of ~30 clusters), the anomaly amplitudes are small (≈0.4 °C standard deviation), and the same sequences appear in four sampling schemes, all of which inflate the apparent precision. The interaction should be re-examined when additional Southern-Hemisphere and equatorial datasets are available. With the present sampling, it does not change the overall conclusion that the temporal increase in episodic positive selection cannot be attributed to temperature.

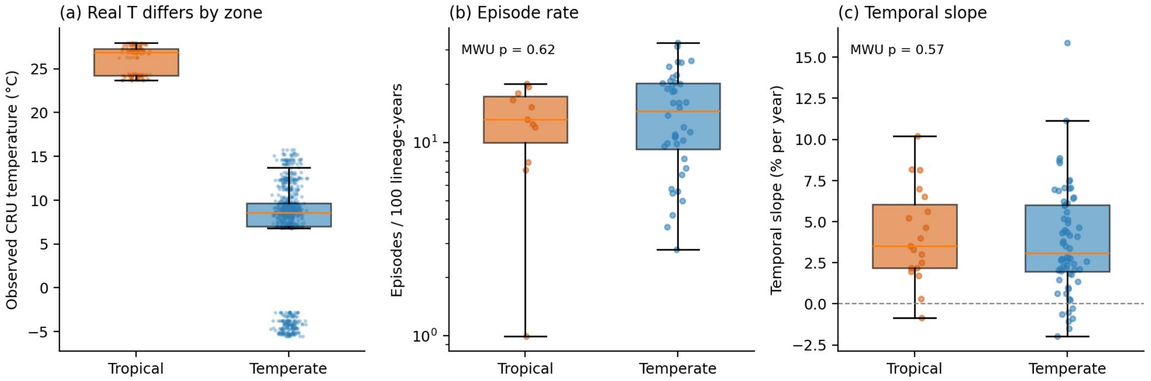


Figure S5.5. Tropical versus temperate, using observed CRU CY v4.08 temperatures. (a) Bin-level observed annual mean temperature differs between zones by ≈20 °C, as expected. (b) Per-dataset episode rate (per 100 lineage-years; pooled across the four sampling schemes; log scale) does not differ between zones (Mann–Whitney $p$ = 0.62). (c) Per-dataset normalised temporal slope (% per year) does not differ between zones (Mann–Whitney $p$ = 0.57). Boxplots show the median and inter-quartile range; jittered points are individual datasets. Orange = tropical (Brazil, India, Singapore, Thailand); blue = temperate (the remaining eight countries).

***Table S6. Per-dataset results of the normalized and raw temporal slopes (CD-HIT-99% scheme). "Fit = no" indicates fewer than 8 dated episodes (or unreliable dates). These datasets contribute to the pooled analysis but were not fit individually.***

| Dataset | n episodes | Norm. slope/yr | Norm. p | Raw slope/yr | Fit |
|---|---|---|---|---|---|
| H1N1 HA china | 29 | +0.0245 | 0.077 | +0.0600 | yes |
| H1N1 HA india | 11 | +0.0217 | 0.489 | +0.0586 | yes |
| H1N1 HA japan | 12 | +0.0214 | 0.463 | +0.0537 | yes |
| H1N1 HA russia | 13 | +0.0520 | 0.073 | +0.0726 | yes |
| H1N1 HA usa | 41 | +0.0200 | 0.004 | +0.0403 | yes |
| H1N1 NA china | 7 | — | — | — | no |
| H1N1 NA india | 23 | +0.0560 | 0.033 | +0.0772 | yes |
| H1N1 NA japan | 3 | — | — | — | no |
| H1N1 NA russia | 5 | — | — | — | no |
| H1N1 NA usa | 13 | +0.0858 | 0.008 | +0.0917 | yes |
| H3N2 HA chile | 9 | +0.0753 | 0.204 | +0.0754 | yes |
| H3N2 HA china | 29 | +0.0235 | 0.204 | +0.0569 | yes |
| H3N2 HA italy | 18 | +0.0707 | 0.027 | +0.0906 | yes |
| H3N2 HA japan | 57 | +0.0261 | 0.030 | +0.0722 | yes |
| H3N2 HA netherlands | 47 | +0.0208 | 0.048 | +0.0540 | yes |
| H3N2 HA russia | 40 | +0.0411 | 0.017 | +0.0828 | yes |
| H3N2 HA singapore | 30 | +0.0030 | 0.872 | +0.0567 | yes |
| H3N2 HA thailand | 39 | — | — | — | excl. |
| H3N2 HA usa | 89 | +0.0185 | 0.083 | +0.0684 | yes |
| H3N2 NA brazil | 4 | — | — | — | no |
| H3N2 NA canada | 3 | — | — | — | no |
| H3N2 NA china | 8 | +0.0696 | 0.095 | +0.0812 | yes |
| H3N2 NA india | 6 | — | — | — | no |
| H3N2 NA italy | 2 | — | — | — | no |
| H3N2 NA japan | 92 | +0.0599 | 1.2×10^-6 | +0.0868 | yes |
| H3N2 NA netherlands | 13 | −0.0054 | 0.730 | +0.0263 | yes |
| H3N2 NA russia | 12 | +0.0278 | 0.236 | +0.0628 | yes |
| H3N2 NA singapore | 12 | +0.1018 | 0.044 | +0.1303 | yes |
| H3N2 NA thailand | 4 | — | — | — | no |
| H3N2 NA usa | 18 | −0.0149 | 0.390 | +0.0395 | yes |

### 7. Supplementary results: which sites are repeatedly under episodic selection

To complement the temporal and climate analyses, we asked where on the protein the selection signal concentrates. For each gene × subtype combination we counted, for every codon site, how many independent episodes (branch × site pairs at MEME [1] $p \leq 0.05$ and EBF > 100) occur there across the CD-HIT-99.7% datasets (the richest of the four schemes), and recorded in how many countries each site is hit.

Selection is strongly concentrated on a small minority of sites in every gene × subtype combination. The top ten sites carry 64.8%, 65.8%, 71.4% and 54.9% of all episodes in HA-H1N1, HA-H3N2, NA-H1N1 and NA-H3N2 respectively; the Gini coefficient of the per-site episode-count distribution is 0.50–0.61 (figure S7, table S7). This is consistent with the well-established observation that positive selection in influenza A surface antigens is focused on a few antigenic and receptor-binding residues [13,14]. Cross-country recurrence is most pronounced for H3N2 HA: 9 sites are under selection in three or more countries, and 3 sites in five or more (figure S7b), most of which pe(figure S7c,d).

The temporal distribution of episodes within each lane of figure S7 recapitulates the through-time acceleration documented in sections §4-6 above: episodes accumulate towards recent decades in essentially every recurrent site, while the spread across countries within a lane illustrates that the antigenic-site signal is globally shared, rather than country-specific. The same sites are being repeatedly targeted by selection in viruses circulating on different continents, year after year, which reflects the molecular signature of ongoing immune escape on a few constrained residues.

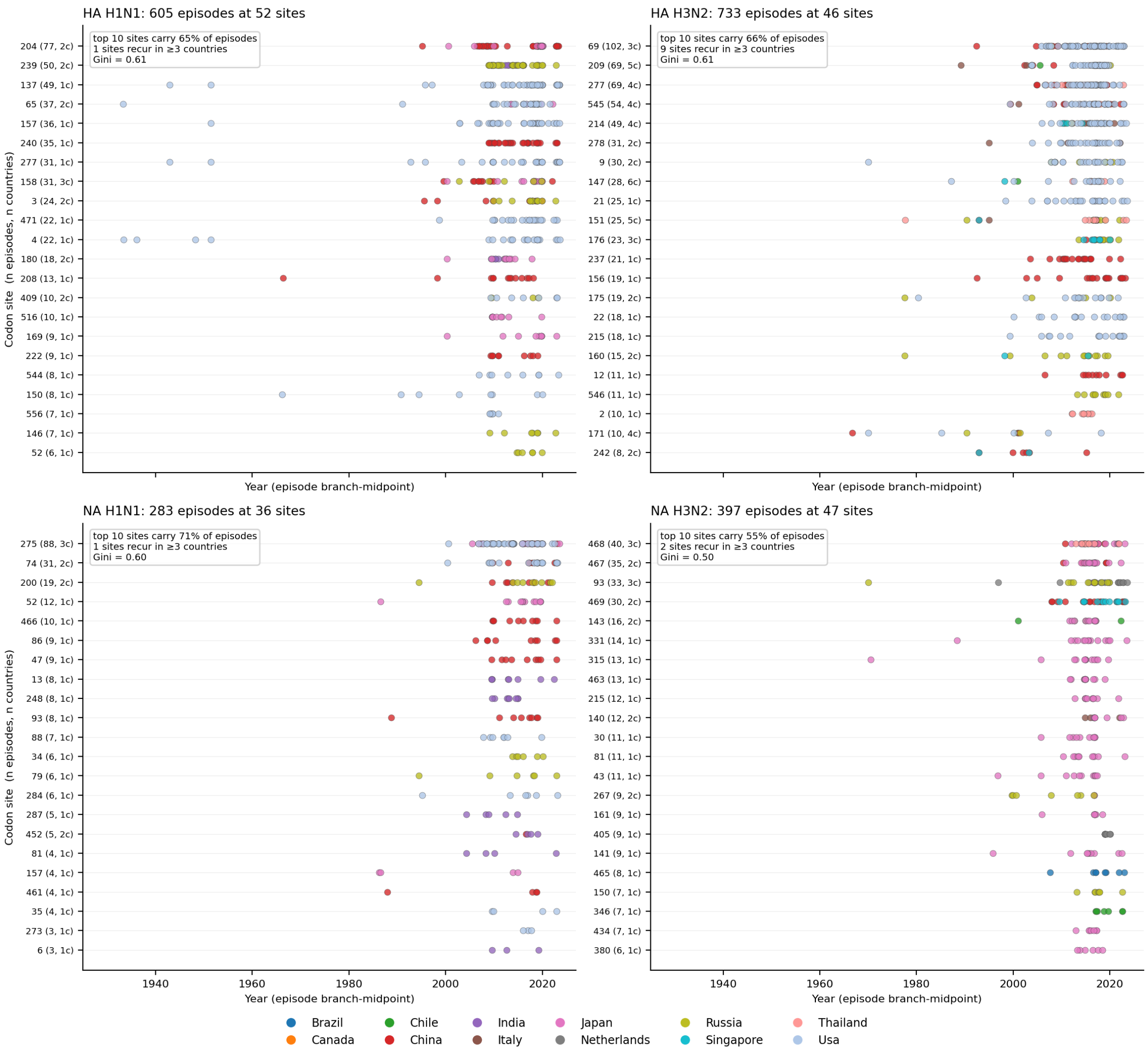


*Figure S7. Codon sites under repeated episodic positive selection, across countries and through time. Each panel shows one gene × subtype combination (CD-HIT-99.7% scheme). Within a panel, each horizontal lane corresponds to one codon site; lanes are ordered by total episode count (most-recurrent at top), and the y-axis label gives the site number, its total number of episodes and the number of countries in which the site is hit (e.g. "145 (32, 6c)" = site 145 with 32 episodes in six countries). Each dot is one episode, placed at its branch-midpoint date and coloured by the country in which the dataset was sampled. Inset boxes give the share of episodes carried by the top ten sites, the number of sites under selection in ≥ 3 countries, and the Gini coefficient of the per-site episode-count distribution. Episodes pile up towards the present and are concentrated on a few sites, most prominently in H3N2 HA.*

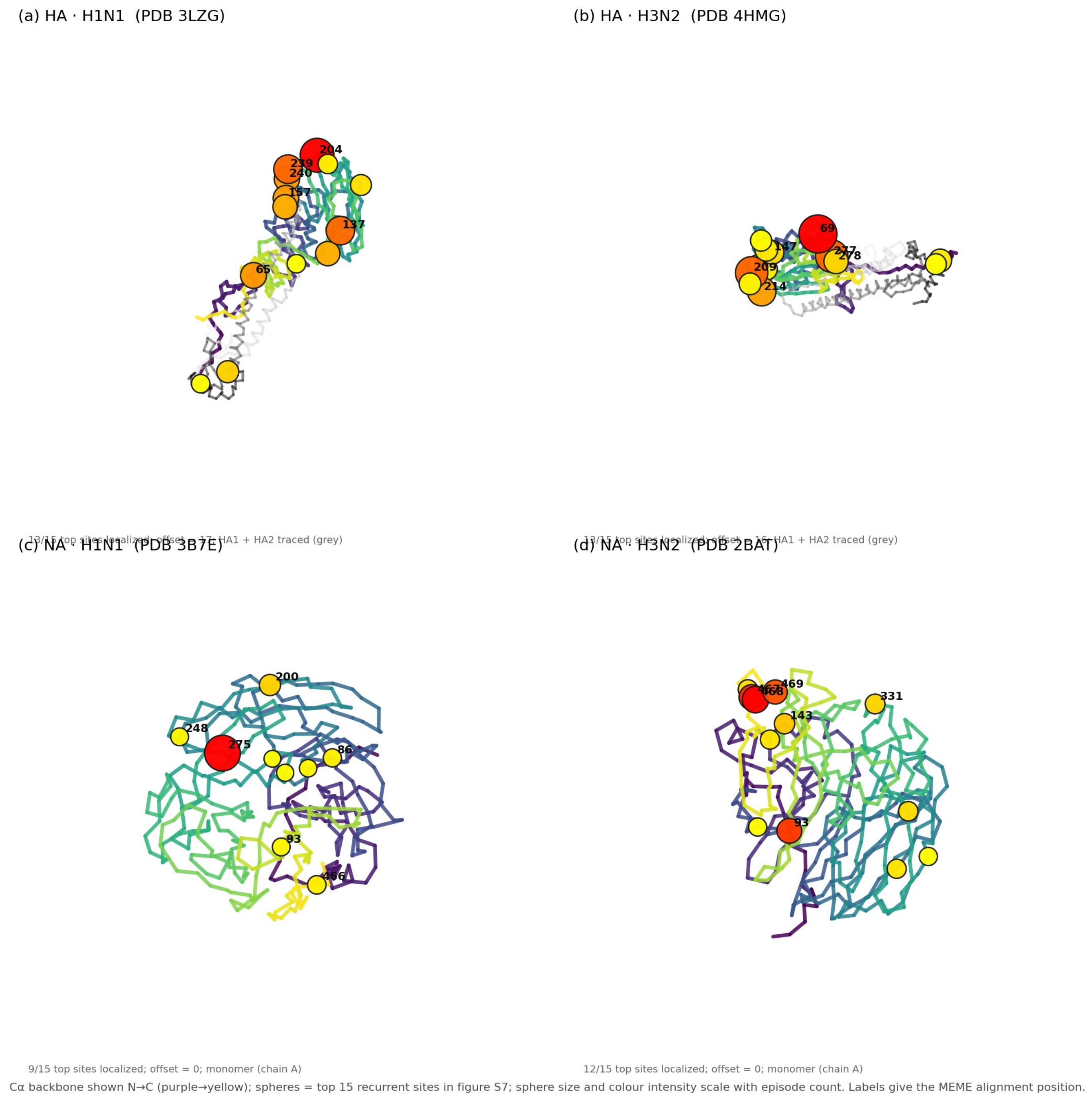


*Figure S8. The top 15 recurrent codon sites from Figure S7 mapped onto representative crystal structures, one panel per gene × subtype. The structures used are: (a) HA · H1N1: PDB 3LZG (2009 pandemic H1 HA), HA1 in viridis, HA2 in grey, signal-peptide offset 17. (b) HA · H3N2: PDB 4HMG (1968 Hong Kong H3 HA, the Wiley/Wilson/Skehel structure), HA1 in viridis, HA2 in grey, offset 16; (c) NA · H1N1: PDB 3B7E (2009 H1N1 NA monomer head), full-length numbering, no offset; (d) NA · H3N2 : PDB 2BAT (1968 Hong Kong N2 NA), full-length numbering, no offset. The Cα backbone is rendered N→C in purple→yellow as a structural guide; the top 15 recurrent sites are drawn as spheres whose size and red intensity scale with episode count, with the six most recurrent labelled by their MEME alignment position. In HA-H3N2 the labelled sites (147 → mature HA1 residue 131; 151 → 135; 156 → 140; 175 → 159; 209 → 193; 277 → 261)*

*cluster at the top of the HA1 head, antigenic sites A, B and D of Wiley, Wilson & Skehel. In HA-H1N1 the largest spheres again sit on the HA1 head (positions corresponding to Sa/Sb/Ca after subtracting the 17-residue signal peptide), consistent with the Caton et al. H1 antigenic-site map. In NA-H1N1 the dominant red sphere is residue 275, the famous H275Y oseltamivir-resistance position; the cluster around residue 200 lies near the catalytic rim. In NA-H3N2 the prominent 467/468/469 cluster sits at the C-terminal end of the head, and 331 falls in the canonical 328-334 antigenic region.*

***Table S7.*** *Concentration and cross-country recurrence of episodic positive selection per gene × subtype. Episodes are matched branch × site pairs at MEME p ≤ 0.05 and EBF > 100 from the CD-HIT-99.7% scheme; the Gini coefficient is computed on the per-site episode-count distribution (0 = uniform, 1 = all episodes at one site).*

| Gene × subtype | Episodes | Unique sites | Top 10 sites (% of episodes) | Sites in ≥ 3 countries | Sites in ≥ 5 countries | Gini |
|---|---|---|---|---|---|---|
| HA × H1N1 | 605 | 52 | 64.8% | 1 | 0 | 0.61 |
| HA × H3N2 | 733 | 46 | 65.8% | 9 | 3 | 0.61 |
| NA × H1N1 | 283 | 36 | 71.4% | 1 | 0 | 0.60 |
| NA × H3N2 | 397 | 47 | 54.9% | 2 | 0 | 0.50 |
